# Rapid generation of conditional knockout mice using the CRISPR-CAS9 system and electroporation for neuroscience research

**DOI:** 10.1101/2021.05.09.443330

**Authors:** Hirofumi Nishizono, Yuki Hayano, Yoshihisa Nakahata, Yasuhito Ishigaki, Ryohei Yasuda

**Affiliations:** Research Support Center, Kanazawa Medical University, 1-1 Daigaku, Uchinada, Kahoku, Ishikawa 920-0293, Japan; Max Planck Florida Institute for Neuroscience, 1 Max Planck Way, Jupiter, Florida 33458, USA

**Author notes:** These authors contributed equally (Hirofumi Nishizono, Yuki Hayano). Correspondence: Hirofumi Nishizono Kanazawa Medical University, 1-1 Daigaku, Uchinada, Kahoku, Ishikawa 920-0293, Japan, Ryohei Yasuda Max Planck Florida Institute for Neuroscience, 1 Max Planck Way, Jupiter, FL 33458, USA.

**Keywords:** Cre/Loxp, floxed mouse, CRISPR-CAS9, genome editing, CaMK1

## Abstract

The Cre/Loxp-based conditional knockout technology is a powerful tool for gene function analyses by allowing region-time-specific gene manipulation. However, inserting a pair of LoxP cassettes for generating conditional knock-out can be technically challenging and thus time- and resource-consuming. This study proposes an efficient, low-cost method to generate floxed mice using the in vitro fertilization and the CRISPR-Cas9 system over two consecutive generations. This method allowed us to produce floxed mice targeting exon 5 to exon 6 of CaMK1 in a short period, 125 days, using only 16 mice. The efficiency of generating floxed mice was 10%, significantly higher than the conventional ES cell-based method. We directly edited the genome of C57BL/6N fertilized eggs, our target genetic background, to eliminate additional backcrossing steps. We confirmed that the genome of this floxed mouse is responsive to Cre protein. This low-cost, highly efficient method for generating conditional knock-out will facilitate comprehensive, tissue-specific genome analyses.

## Main Text

The invention of ES cell-based gene targeting technology in mice and conditional knockout systems using Cre/Loxp have led to breakthroughs in neuroscience [1, 2]. These technologies have allowed researchers to analyze gene function in a region-specific or time-specific manner. However, in the past, the generation of gene-modified mice required a long time and advanced technology [3, 4]. Injection of DNA donor into a fertilized egg's pronucleus using microinjection required highly skilled personnel and expensive equipment and had suffered from a low success rate. Furthermore, the requirement of backcrossing after the generation of chimeric mice further slows the production cycle. Recently, the development of genome editing technology for fertilized eggs combined with electroporation and the CRISPR-Cas9 system (TAKE and iGONAD) has made it possible to generate gene-modified mice easily and with a high success rate using inexpensive equipment [5–7]. However, electroporation permits only short (1kb or less) single-stranded DNA donors to be introduced into the nucleus [8]. Thus, while the electroporation-based method allows for the efficient development of the knock-in of a relatively short tag, it is not suitable for creating conditional knockout mice for the deletion of a long genomic region such as LoxP- multiple exons-LoxP cassettes.

A research group has recently proposed solutions to the problem of DNA donor size limitation [9]. This method, Easi-CRISPR, can efficiently insert relatively long transgene into the genome of mouse fertilized eggs using long single-strand DNA (ssDNA) as a DNA donor. The use of long synthesized long ssDNA and CRISPR ribonucleoproteins (ctRNPs) is a unique feature of this method. However, this approach does not allow researchers to generate floxed mice for deleting regions larger than 1.5 kb. Another problem is that long ssDNA is often challenging and expensive to synthesize. Other research groups have reported that it is possible to introduce LoxP sequences on the 5’ and 3’ sides by sequential electroporation two times during the fertilized egg’s embryonic development [10, 11]. This method could theoretically be used even if the region to be knocked out is more than 1.5 kb using only electroporation. However, it has been difficult to reproduce this method in a large-scale replication study with multiple experimental facilities [12]. We also conducted a replication study, but the results were the same (data not shown).

As a compromise between the simplicity and the time-cost of generating conditional knockout mice with more than 1.5 kb, we tried to generate conditional knockout mice through two generations (Fig 1. A and B). Exon 5 and exon 6 of Ca^2+^/calmodulin-dependent protein kinase 1 (CaMK1) [13] were designed to be knocked out under Cre recombinant expression. Briefly, a DNA donor containing a LoxP and a homology arm (5’ LoxP) is knocked into the upstream intron of exon 5. A similar short construct is also inserted into the downstream intron of exon 6 (3’ LoxP). These two DNA donors are single-stranded oligo donors (ssODNs) of less than 200 bases each, and thus they can be transferred into the pronucleus of the fertilized eggs by electroporation. First, we knocked-in 5’ LoxP into C57BL/6 mouse embryos and then transferred the embryos to pseudopregnant females. When the resulting 5’ LoxP-bearing mice are 6 to 8 weeks old, we performed a second round of in vitro fertilization (IVF) using their sperm with wild type oocytes. The fertilized eggs carrying 5’ LoxP were then electroporated with a 3’ LoxP DNA donor together with gRNA and Cas9 protein. The electroporation was performed as described in our previous paper [14]. As a result, four mice carrying both 5’ and 3’ LoxP were born among ten mice in the second round of IVF. Note that when introducing 5’ and 3’ LoxP separately in two rounds of genome editing, it is necessary to confirm that both DNA donors are inserted into the same allele. Therefore, the long PCR in Fig. 1C was performed using PCR primers, including both LoxP sites (Fig. 1A). This result shows that both LoxPs are introduced into the same allele in one of the mice. The sequencing results indicate that LoxP is introduced in the correct position and direction (Fig. 1D).

**Figure 1.**
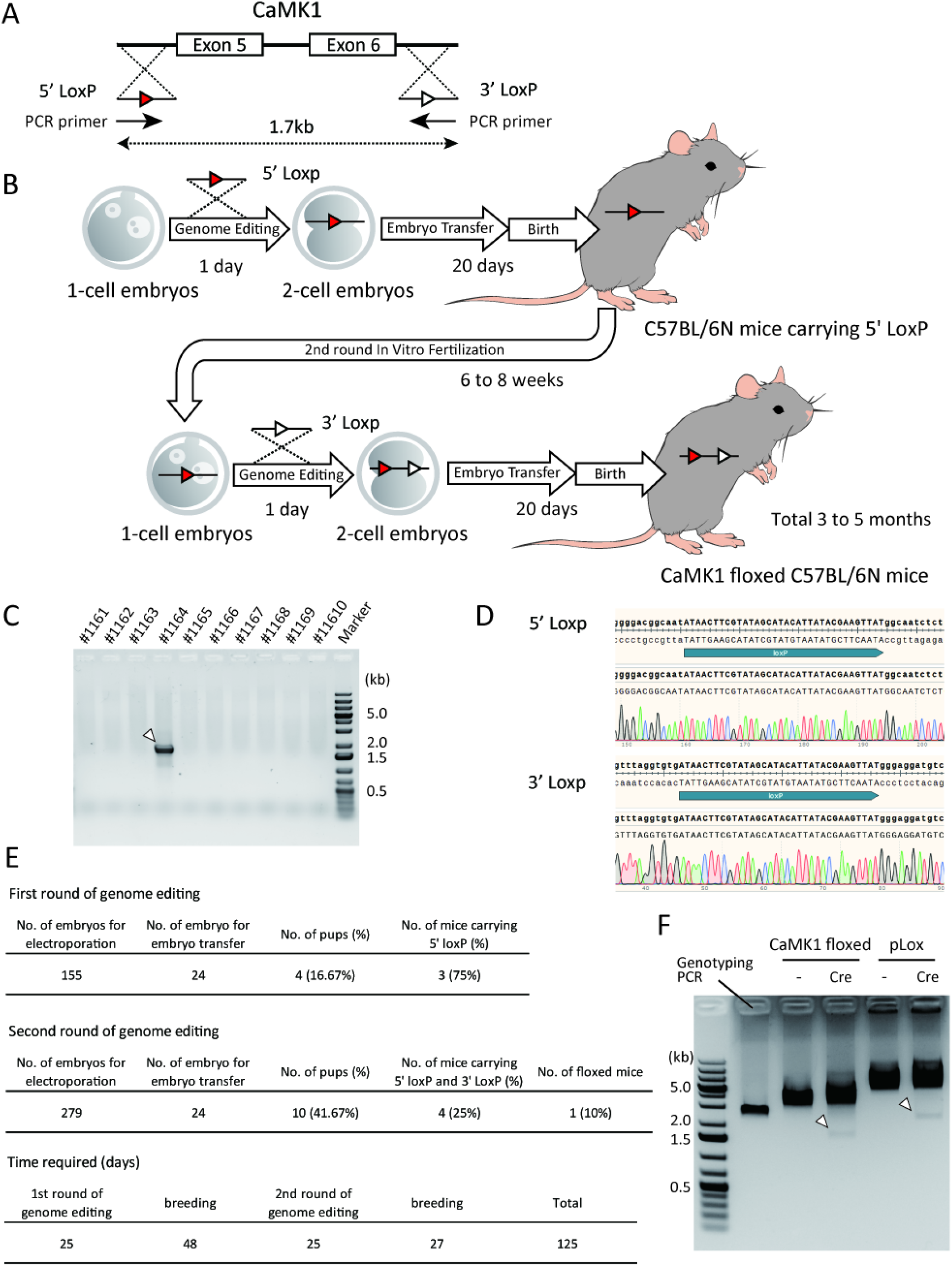
Generation of CaMK1 floxed mice. (A) Diagram of the constructs. gRNAs and ssODNs were designed to introduce a 5’-side LoxP (5’ LoxP) into the intron upstream of Exon 5 and a 3’-side LoxP (3’ LoxP) into the intron downstream of Exon 6. The distance between the Loxps is 1.7 kb. (B) Scheme of sequential in vitro fertilization and genome editing over two generations. (C) PCR results of ten mice born after two rounds of genome editing. Arrowheads indicate target bands. (D) DNA sequencing around LoxP of CaMK1 floxed mouse (#1164). The upper panel is around 5’ LoxP, and the lower panel is around 3’ LoxP. No deletion site or mismatch could be identified. (E) Detailed results of the first and second rounds of genome editing. (F) Results of Cre protein treatment of the PCR product from CaMK1 floxed mouse genome. Arrowheads indicate the excised DNA. Cre; Cre protein treated group, -; Cre protein non-transfected group pLOX; control DNA (pLOX) was used instead of the PCR product from CaMK1 floxed mouse genome.

The efficiency of this method for generating floxed mice is summerized in Fig. 1E. The first round of IVF and electroporation produced four mice, three of which had 5’ LoxP. In the second round of genome editing, ten mice were born, and 4 of them had both 5’ and 3’ LoxP. Of these, one was a floxed mouse that had both Loxps in the same allele. There was no mouse with 3’ LoxP alone. These results indicate that each genome editing treatment was efficiently introduced ssODN into the genome. The second genome editing produced floxed mice with a 10% success rate, much more efficient than previous DNA microinjection-based methods (2 to 5%) [3, 15]. The number of wild-type C57BL/6N female mice used in these experiments was 7 in the first round and 9 in the second round. Thus, we succeeded in creating a new floxed line with fewer than 20 mice using this method. Theoretically, the time required could be as short as three months, but it took 125 days in practice, as shown in Fig. 1C. Even so, it is possible to produce the desired floxed mice in a remarkably short time. When the CaMK1 floxed mouse genome was treated with recombinant Cre protein in a cell-free assay, the electrophoresis pattern was changed only in the presence of Cre protein, as shown in Fig. 1F. This result indicates that the genome of the generated CaMk1 floxed mice can induce a conditional knockout with Cre protein. We genome-edited C57BL/6N fertilized eggs, the genetic background of interest, so no backcrossing was required. For some sensitive assay, backcrossing with wild-type C57BL/6N may be necessary to eliminate potential off-target genome editing.

We have shown that it is possible to generate new floxed mice from as few as 16 mice in as little as 125 days. While how general the described approach is for other genes and species required to be followed up, we believe that our method will provide a highly efficient and cost-effective means to develop new conditional knock-in mice for many different genes in the future.

## Materials and Methods

### Animals

Mice aged 8 to 12 weeks with the target genetic background, C57BL/6N, were used for the experiment (Charles River, Wilmington, MA). Animals were maintained under a 12:12-h light: dark cycle at 22 ± 2°C and relative humidity of 40–60% with access to chow and water *ad libitum*.

### Genome Editing

Genome editing was performed using the same method as in our previous paper. Our modification for this study was the use of fresh in vitro fertilized eggs instead of frozen embryos. Two crRNAs (5’-tatgcaccaggggacggcaatgg-3’ UAUGCACCAGGGGACGGCAAGUUUUAGAGCUAUGCU for 5’ Loxp, 5’-ggtgtgatccggtttaggtgtgg −3’ GGUGUGAUCCGGUUUAGGUGGUUUUAGAGCUAUGCUfor 3’ Loxp) and one tracrRNA were used in the experiment. Two types of ssODNs, 5’-ctgcacgacctgggcattgtgcaccgggatctcaaggtaggatctgaggggcctagtgaactatatgcaccaggggacggcaatATAACTTCGTATAGCATACATTATACGAAGTTATggcaatctctgtctgtcctgctttgtctgtctttgagtacctctcagcccctcactaaagccctagctttccatttgcaa-3’ and 5’-tcacttcagatagtcaaaggccctttgtgatggtaaaatctgagtggcttttgagccagtttaggtgtgatccggtttaggtgtgAT AACTTCGTATAGCATACATTATACGAAGTTATgggaggatgtcaaacatgaagaccctatgacagcatgttcaag gacagaaggaaggccagtactgccagacagaagtgag-3’, were used as a transgene, respectively. We also used 5’-acattatacgaagttatggcaatct-3’ and 5’-gctatacgaagttatcacacctaaacc-3’ primers for genotyping PCR. All DNA and RNA oligos were purchased from Integrated DNA Technologies, Inc (Coralville, IA).

### Cre protein treatment

Long PCR was performed using the primers 5’-ggtttcaggtggagagctgt-3’ and 5’-cagagtcagagatgtcgtccca-3’ for the genome of the CaMK1 floxed mice. The Purified PCR product was treated with Cre recombinase as described in the instructions (M0298, New England Biolabs, Ipswich, MA). The CaMK1 floxed PCR product and control plasmid (pLOX) were incubated at 37°C for 30 minutes with or without the Cre protein. Then the Cre was denatured at 70°C for 10 minutes. DNA cleavage was confirmed by electrophoresis.

## Declarations

### Ethics approval and consent to participate

All animal experiments were conducted according to the guidelines and the rules of the Institutional Animal Care and Use Committee of Max Planck Florida Institute for Neuroscience. The experimental protocol was approved (Approval Number: 19-001).

### Consent for publication

Not applicable.

### Availability of data and materials

The authors declare that all data supporting the findings of this study are available within the article.

### Competing interests

The authors declare that they have no competing interests.

### Funding

This study was funded by NIH (R01MH080047, R35NS116804).

### Authors’ contributions

H.N. and R.Y. designed the studies and performed the analysis of the results. H.N. and Y.H. were responsible for the actual genome-edited knock-in mouse generation and genome analysis. Y.N. designed the knock-in constructs and did the experiments. Y.I. analyzed the experimental results and revised the manuscript. All authors read and approved the final manuscript.

## Acknowledgements

We would like to thank Amanda Coldwell, Elizabeth Garcia, and Idris El-Amin for the care and management of the animals.

## Notes

### Competing Interest Statement

The authors have declared no competing interest.

